# Comparison of Human Intestinal Enteroids and Zebrafish Larvae models for replication of Human Norovirus

**DOI:** 10.64898/2026.01.21.700872

**Authors:** Sahaana Chandran, Jan Vinjé, Kimberly Huynh, Kristen E. Gibson

## Abstract

Human intestinal enteroids (HIE) and zebrafish larvae (ZF) models support the replication of certain strains of human norovirus (HuNoV), the leading cause of viral gastroenteritis worldwide. The replication of 17 HuNoV (9 genotypes) positive stool specimens from patients ranging from 5 months to 83 years old was evaluated in both HIE and ZF models. The yolk of 3-day post-fertilization ZF larvae were microinjected with 3 nL 10% clarified stool suspension. Each day post-infection (dpi), 10 larvae were pooled as one sample, and two specimens were collected daily until 5 dpi. Viral RNA was extracted from harvested larvae and quantified using reverse transcription (RT) droplet digital PCR. For the HIE model, J4FUT2 K1 enteroid monolayers were inoculated with 100-fold dilution of each stool specimen. Viral RNA levels were quantified at 1 and 72 h post-infection using RT real-time PCR. All genogroup (G) I specimens (n=5) replicated in both models showing a 1 to 2.5 log increase in HuNoV RNA. Of the 12 GII specimens tested, 8 replicated in the ZF model and 7 replicated in the HIE model. Specimens that failed to replicate in each model were not the same, although most of them were GII.4 Sydney [P16]. One additional strain, GII.17[P17], replicated only in the ZF model. Among the GII strains, viral RNA increase ranged from 1.6 to 3.3 log in the ZF and 0.4 to 3.68 log in the HIE model. These data highlight variability in HuNoV replication efficiency across models, potentially indicating the influence of model-specific factors on replication.

**Importance:** Human intestinal enteroid and ZF models have been independently shown to support replication of different strains of HuNoV, the leading cause of viral gastroenteritis worldwide. However, whether the same HuNoV positive stool specimens replicate similarly in both systems has not been examined. In this study, stool specimens positive for a range of HuNoV genotypes were tested in both models. While specimens from the GI genogroup consistently replicated in both systems, specimens from the GII genogroup showed variable replication, with some replicating in only one model and a couple of specimens not replicating in either model. These findings demonstrate that model specific biological factors influence HuNoV replication and highlight the significance of using complementary model systems to better understand HuNoV pathogenesis.

## Introduction

Human norovirus (HuNoV) is the leading cause of viral gastroenteritis globally, contributing to an estimated 136,000 to 278,000 deaths each year (1). The annual global economic burden of HuNoV is substantial, with direct healthcare costs estimated at $4.2 billion and societal costs reaching $60.3 billion (2). Despite its significant impact, there are currently no approved vaccines or antiviral treatments available for HuNoV (3), underscoring the urgent need to better understand its pathogenesis and to develop effective intervention strategies. Human norovirus research has been significantly hampered due to the lack of a reliable cell culture model that supports the replication of HuNoV (4).

The advent of the human intestinal enteroid (HIE) model in the year 2016 marked a breakthrough in HuNoV research (5). Derived from stem cells isolated from the intestinal crypts of the human intestinal tissue, these cells can recapitulate the cellular diversity and physiology of the human intestinal epithelium (5). Human intestinal enteroids have supported the replication of multiple GI and GII HuNoV strains, providing a valuable tool for studying strain specific differences and host interactions (5, 6, 7, 8, 9, 10, 11). Recently, it has been shown that in genetically modified HIE lines (J4FUT2-KI and J8FUT2-KI), GII.3 strain can be passaged for 10-15 rounds in the presence of TAK-779, an antagonist of chemokine receptors (CXCR3, CCR5, CCR2) (12).

Another model available for studying HuNoV is the zebrafish model. Both zebrafish embryos and larvae (at 3 days post fertilization [dpf]) have been shown to support the replication of multiple GII HuNoV strains through a microinjection method (13, 14, 15). Furthermore, zebrafish larvae have been shown to support continuous passaging of HuNoV strain GII.4 up to two cycles (13), and zebrafish embryos reportedly support passaging up to four times for three different GII HuNoV strains (GII.2, GII.4, GII.17) (14).

Although both the HIE model and the zebrafish larvae or embryo model have been shown to support the replication of various HuNoV strains, with some strains replicating in both HIE and zebrafish model, no prior study has directly compared the replication of the same stool specimens in both models. In this study, we evaluated five GI and 12 GII stool specimens to determine replication across these models and if differences in replication outcomes arise due to the distinct biological environments each model presents.

## Materials and Methods

### Laboratory sites

The study was conducted jointly by the National Calicivirus Laboratory at the U.S. Centers for Disease Control and Prevention (CDC; Atlanta, GA) for the HIE model and by the University of Arkansas System Division of Agriculture (Fayetteville, AR) for the zebrafish model.

### Human norovirus stool samples

Seventeen HuNoV positive stool specimens were obtained from outbreaks and sporadic cases of acute gastroenteritis. **Table 1** lists the specific HuNoV genotypes as well as the age of the patient obtained from and initial viral RNA titers.

**Table 1:** Human norovirus strains used in this study.

| Human Norovirus<br>Genotype | Patient Age | Titer (RNA copies/g of stool) |  |
| --- | --- | --- | --- |
|  |  | ddPCR<sup>a</sup> | RT-qPCR |
| GI Genogroup |  |  |  |
| GI.3 [P3] | 71 y | 1.43 x 10<sup>11</sup> | 1.00 x 10<sup>11</sup> |
| GI.5 [P5] a | 69 y | 2.15 x 10<sup>10</sup> | 4.77 x 10<sup>7</sup> |
| GI.5 [P5] b | 80 y | 5.84 x 10<sup>9</sup> | 1.54 x 10<sup>7</sup> |
| GI.5 [P5] c | 67 y | 1.64 x 10<sup>9</sup> | 1.84 x 10<sup>9</sup> |
| GI.7 [P7] | 81 y | 3.22 x 10<sup>10</sup> | 1.92 x 10<sup>11</sup> |
| GII Genogroup |  |  |  |
| GII.3 [P12] a | 5 m | 4.48 x 10<sup>9</sup> | 9.12 x 10<sup>9</sup> |
| GII.3 [P12] b | 68 y | 4.56 x 10<sup>9</sup> | 3.89 x 10<sup>9</sup> |
| GII.4 Sydney [P16] a | 5 m | 4.83 x 10<sup>9</sup> | 4.88 x 10<sup>9</sup> |
| GII.4 Sydney [P16] b | 13 m | 4.86 x 10<sup>9</sup> | 6.02 x 10<sup>9</sup> |
| GII.4 Sydney [P16] c | 83 y | 3.44 x 10<sup>9</sup> | 3.03 x 10<sup>9</sup> |
| GII.4 Sydney [P16] d | 69 y | 8.90 x 10<sup>9</sup> | 6.53 x 10<sup>9</sup> |
| GII.4 Sydney [P16] e | 4 m | 3.21 x 10<sup>9</sup> | 2.39 x 10<sup>9</sup> |
| GII.4 Sydney [P16] f | 89 y | 3.83 x 10<sup>9</sup> | 5.09 x 10<sup>9</sup> |
| GII.4 Sydney [P31] | 5 m | 3.07 x 10<sup>10</sup> | 2.06 x 10<sup>9</sup> |
| GII.4 [P4 New Orleans] | 16 y | 1.11 x 10<sup>9</sup> | 2.70 x 10<sup>8</sup> |
| GII.6 [P7] | 1 y | 8.26 x 10<sup>9</sup> | 2.55 x 10<sup>10</sup> |
| GII.17 [P17] | 14 y | 4.40 x 10<sup>8</sup> | 6.89 x 10<sup>8</sup> |

<sup>a</sup> ddPCR – Droplet Digital Polymerase Chain Reaction
<sup>b</sup> RT-qPCR: Reverse Transcription-quantitative Polymerase Chain Reaction

### Human intestinal enteroid model for HuNoV infection

#### Human intestinal enteroid culture

The J4FUT2-KI cell line, genetically-modified in which the *fucosyltransferase* 2 gene was knocked into parental J4 HIEs using CRISPR-Cas9 (Haga et al., 2020) (provided by Dr. Mary Estes, Baylor College of Medicine, Houston, TX), was grown as 3D cultures in Matrigel (BD Biosciences, San Diego, CA, USA) with IntestiCult™ Organoid Growth Medium Human Basal Medium (STEMCELL Technologies, Seattle, WA, USA) with 10 μmol/L ROCK inhibitor (Sigma-Aldrich, St. Louis, MO, USA) as described (6).

HIE monolayers were prepared from 3D HIEs and seeded into collagen-coated 96-well plates as described (6). After 24 h, culture medium was replaced with differentiation medium to induce cell differentiation. Differentiation media was composed of equal volumes of complete medium without growth factors [CMGF-; Advanced Dulbecco Modified Eagle Medium (DMEM)/F12 (Invitrogen) supplemented with 1% GlutaMAX (Invitrogen), 1% Penicillin/Streptomycin (Invitrogen), and 10 mM HEPES (Invitrogen)] and IntestiCult® OGM basal medium (STEMCELL Technologies). Cells were differentiated for 5 days. Media was refreshed every other day.

#### Preparation of 10% clarified stool samples

Ten percent HuNoV-positive stool filtrates were prepared as described previously (6). Briefly, 4.5 g of ice-cold 1× phosphate-buffered saline (PBS) was added to 0.5 g of stool, homogenized by vortexing for 1 min. The suspension was then centrifuged at 1,500 × g for 10 min at 4°C. The supernatant was transferred to a new tube and stored in aliquots at −80°C until use.

#### HuNoV infection of HIE

Clarified stool specimens were thawed and centrifuged at 10,000 x g for 10 min. The supernatant was diluted 1/100 in CMGF(-) and 100 μL was inoculated on 5-day old differentiated HIE monolayers and incubated at 37°C and 5% CO_2_. After 1 h post-infection, 96-well plates were washed twice with CMGF(-) to remove unbound viruses. Differentiation medium (100 μL/well) supplemented with 500 μM glycochenodeoxycholic acid [GCDCA] (Sigma-Aldrich) plus 50 μM ceramide (Santa Cruz Biotechnology, Dallas, Texas, USA) was then added to the HIEs and incubated at 37°C and 5% CO_2_. One plate was immediately frozen at -70°C, and a duplicate plate was incubated at 37°C, 5% CO_2_ for 72 h and frozen at -70°C.

#### RNA Extraction and reverse transcription real-time PCR (RT-qPCR)

Norovirus RNA was extracted from the inoculated HIE cultures using the KingFisher Flex (Thermo Fisher Scientific) and the MagMAX-96 Viral Isolation Kit (Thermo Fisher Scientific) following the manufacturer’s specifications. Norovirus GI/GII RT-qPCR was used to determine the amount of HuNoV RNA present in the inoculum and infected cells (6). A 10-fold serial dilution of a quantified GI.4 and GII.4 Sydney norovirus RNA transcript was included in each assay to generate a standard curve for RNA quantification. Viral replication was quantified by measuring the increase in virus genome equivalents at 72 h compared to 1 h post-infection.

### Zebrafish model for HuNoV infection

#### Zebrafish Husbandry and Maintenance

Adult wild-type zebrafish (AB strain) were obtained from the Zebrafish International Resource Center (ZIRC, Eugene, OR, USA) and were kept in a recirculating tabletop aquaculture system (eRack; Aquaneering, Inc., San Marcos, CA, USA). Fish were housed at a temperature of 28.5°C with a photoperiod of 14 h light and 10 h dark. Fertilized embryos were collected by spawning adult zebrafish in mating tanks and then they were kept in petri dishes with Danieau’s solution (composed of 1.5 mM HEPES, 17.4 mM NaCl, 0.21 mM KCl, 0.12 mM MgSO_4_, 0.18 mM Ca(NO_3_)_2_, and 0.6 μM methylene blue). All experimental procedures involving zebrafish were conducted in accordance with protocols approved by the Institutional Animal Care and Use Committee at the University of Arkansas, Fayetteville (AUP22020, AUP23012).

#### Preparation of HuNoV stool samples for zebrafish embryo and larvae microinjection

Each of the HuNoV stool specimens (Table 1) obtained from the National Calicivirus Laboratory of the CDC was processed by taking 100 mg of stool specimen and suspending it in 1 mL of 1×PBS, followed by vortexing and centrifuging at 9,000 × *g* for 5 min. The supernatant was collected, aliquoted, and stored at -80°C for further use.

#### Microinjection of zebrafish embryos and larvae with HuNoV sample

For microinjection, injection needles were pulled out of glass capillaries using a micropipette puller with a heat filament (Sutter Instruments, Novato, CA, USA). Microinjection was done using a M3301R Manual Micromanipulator (World Precision Instruments, Sarasota, FL, USA) and a Nanoject III Microinjector (Drummond Scientific Company, Broomall, PA, USA). For embryo microinjection, fertilized eggs were collected and rinsed three times using 0.3× Danieau’s solution. The embryos were then transferred to a petri dish molded with six rows of angled grooves in 1.5% agarose (World Precision Instruments). Each zebrafish embryo was injected within 4 h post-fertilization (hpf) with 3 nL of clarified HuNoV stool specimen. Control embryos were injected with 3 nL of 1×PBS. For larvae microinjection, 3 dpf larvae were anesthetized by immersion in Danieau’s solution containing 0.4 mg/mL tricaine (Sigma-Aldrich) for 2 min. The anesthetized larvae were then positioned in a petri dish molded with six rows of angled grooves in 1.5% agarose. Larvae were aligned so that their yolk sacs faced the injection needle. Each larva was then microinjected with 3 nL of clarified HuNoV stool specimen. Control embryos and larvae were injected with 3 nL of 1×PBS. After microinjection, zebrafish embryos or larvae were transferred into 6-well plates, with each well containing 10 mL of 0.3× Danieau’s solution and ten embryos or larvae. The plates were kept in an incubator at 32°C with a 14/10 h light/dark cycle. Post-injection, embryo and larval development was monitored daily, with any dead embryos or larvae being removed and the medium being changed daily. Ten zebrafish embryos or larvae were pooled as one sample, and two samples were collected until 5 days post-infection (dpi) into 2 mL tubes and stored at -80°C for further processing.

#### RNA extraction from zebrafish embryos and larvae

Zebrafish embryos and larvae stored at - 80°C in 2 mL tubes were mixed with 800 μL of TRI reagent (Zymo Research, Irvine, CA, USA) and then homogenized using a handheld VWR 200 Homogenizer Unit fitted with a 5 x 75 mm flat-bottom generator probe (VWR International, Radnor, PA, USA). The homogenizer was set to position 1, corresponding to a speed range of 5,000 to 8,000 rpm. The embryo and larvae samples were homogenized for 3 cycles of 15 s, with rest intervals of 60 s. The homogenates were clarified by centrifugation at 9,000 × *g* for 5 min, and RNA was extracted using the Direct-zol RNA Miniprep (Zymo Research) following the manufacturer’s protocol.

#### Droplet digital PCR for quantification of viral RNA copies in zebrafish samples. Droplet digital

PCR (ddPCR) was used to quantify viral RNA copy numbers. Reactions were prepared using the One-Step RT-ddPCR Advanced Kit for Probes (Bio-Rad Laboratories, Hercules, CA), with a final concentration of 900 nM for each primer, 250 nM for the probe, and 15 nM dithiothreitol (DTT). Primer and probe sequences specific to genogroup (G)I and GII HuNoV strains were as follows: GI forward primer QNIF4 (5′-CGCTGGATGCGNTTCCAT -3′), GI reverse primer NV1LCR (5′-CCTTAGACGCCATCATCATTTAC -3′), GI probe (5FAM/TGGACAGGA/ZEN/GAYCGCRATCT/3IABkFQ), GII forward primer QNIF2 (5′-ATGTTCAGRTGGATGAGRTTCTCWGA-3′), GII reverse primer COG2R (5′-TCGACGCCATCTTCATTCACA-3′), and GII probe (5FAM/AGCACGTGG/ZEN/GAGGGCGATCG/3IABkFQ). All oligonucleotides were purchased from Integrated DNA Technologies, Inc. (San Diego, CA, USA).

Each 20 μL reaction was loaded into DG8™ Cartridges (Bio-Rad Laboratories), followed by the addition of 70 μL of Droplet Generation Oil for Probes (Bio-Rad Laboratories). Droplets were generated using the QX200™ Droplet Generator (Bio-Rad Laboratories), and then 40 μL of the droplet suspension was transferred to a 96 well PCR plate by pipetting. Plates were sealed using the PX1 PCR Plate Sealer (Bio-Rad Laboratories) and then were PCR amplified on a C1000 Touch™ Thermal Cycler (Bio-Rad Laboratories). The cycling conditions were as follows: reverse transcription at 47°C for 60 min, enzyme activation at 95°C for 10 min, followed by 40 amplification cycles of 95°C for 30 sec (denaturation, 3°C/s ramp rate), and 53°C for 1 min (annealing/extension, 3°C/s ramp rate). A final enzyme deactivation step was performed at 98°C for 10 min. PCR plates were held at 4°C overnight prior to analysis on the QX200™ Droplet Digital™ PCR System. Data were processed using QuantaSoft™ version 1.7.4 software, with manual thresholding to distinguish positive from negative droplet clusters.

## Results

### Zebrafish larvae and HIE model for replication of GI strains of HuNoV

Five different GI strain positive human stool specimens were studies using both the HIE and zebrafish larvae model. The strains included GI.3[P3], GI.5[P5] (3 different stool specimens) and GI.7[P7] (**Table 1**). All five stool specimens were from elderly patients (>65 years), and all of them replicated in both the models (**Table 2**). In the zebrafish larvae model, the GI.3[P3] strain showed a maximum log increase of 1.14 at 3 dpi, while the HIE model showed 1.24 log increase at 72 hpi. For the three different GI.5[P5] stool specimens, the zebrafish larvae model showed maximum log increases of 1.98, 1.69 and 2.62 at 1 dpi, and the corresponding HIE model results showed log increases of 2.15, 1.28, and 2.65 after 72 hpi, respectively. With GI.7[P7], a 1.77 log increase was observed on 1 dpi in the zebrafish larvae model, and a 1.15 log increase was detected at 72 hpi with the HIE model.

**Table 2:** Log increase in each human norovirus strain observed in the zebrafish larvae model and HIE model.

| Genogroup | Genotype | Average Log Increase in Viral RNA |  |
| --- | --- | --- | --- |
|  |  | Zebrafish model<br>(per larva)<sup>a</sup> | HIE model |
| GI | GI.3 [P3] | 1.14 | 1.24 |
| GI | GI.5 [P5] a | 1.98 | 2.15 |
| GI | GI.5 [P5] b | 1.69 | 1.28 |
| GI | GI.5 [P5] c | 2.62 | 2.65 |
| GI | GI.7 [P7] | 1.77 | 1.15 |
| GII | GII.3 [P12] a | 2.72 | 2.94 |
| GII | GII.3 [P12] b | NR | NR |
| GII | GII.4 Sydney [P16] a | 1.61 | 0.65 |
| GII | GII.4 Sydney [P16] b | 2.56 | NR |
| GII | GII.4 Sydney [P16] c | 1.49 | NR |
| GII | GII.4 Sydney [P16] d | 3.2 | 3.68 |
| GII | GII.4 Sydney [P16] e | NR | 2.39 |
| GII | GII.4 Sydney [P16] f | NR | 0.42 |
| GII | GII.4 Sydney [P31] | 2.02 | 2.54 |
| GII | GII.4 [P4 New Orleans] | NR | NR |
| GII | GII.6 [P7] | 2.2 | 1.06 |
| GII | GII.17 [P17] | 2.04 | NR |

NR – No replication observed
<sup>a</sup> Two experimental trials were conducted for each stool specimen and two samples were collected for each day. The average log increase was calculated by subtracting the mean log viral RNA copies detected on Day 0 from the mean log viral RNA copies detected on the day showing the maximum increase in viral RNA.

### Zebrafish larvae and HIE model for replication of GII strains of HuNoV

Twelve different GII stool specimens were tested using zebrafish larvae and HIE model. The strains included GII.3[P12] (2 different stool specimens), GII.4 Sydney[P16] (6 different stool specimens), GII.4 Sydney[P31], GII.4[P4 New Orleans], GII.6[P7] and GII.17[P17] (Table 1). Previous studies have reported that HuNoV GII positive stool specimens from pediatric patients (<5 years of age) replicate better in the HIE model compared to stool specimens from older patients (6, 7, 16). In the present study, six GII stool specimens from pediatric patients (<2 years of age), two GII stool specimens from teenage patients (14 years and 16 years), and four stool specimens from elderly patients (>65 years) were tested. In the HIE model, 83% (5 of 6) of pediatric stool specimens replicated successfully, neither of the teenage specimens replicated, and 50% (2 of 4) of elderly stool specimens replicated. With the zebrafish larvae model, no previous study has considered the effect of patient age on successful replication. In the zebrafish larvae model, 83% (5 of 6) of the pediatric stool specimens replicated, 50% (1 of 2) of teenage stool specimens replicated, and 50 % (2 of 4) of stool specimens from elderly patients replicated.

While one GII.3[P12] replicated in both the models, with a maximum log increase of 2.72 at 2 dpi in the zebrafish larvae model and a log increase of 2.94 in HIE model at 72 hpi, the other GII.3[P12] strain did not replicate in either of the models (Table 2). A maximum log increase of was seen on 1 dpi in the zebrafish model for GII.4 Sydney[P31], and in the HIE model the same strain had a 2.54 log increase at 72hpi (Table 2). The strain GII.4[P4 New Orleans] did not replicate in both the models (Table 2). With the strain GII.6[P7], a maximum log increase of 2.20 was seen in the zebrafish larvae model on 2 dpi whereas with the HIE model a 1.06 log increase at 72 hpi was observed (Table 2). Replication of GII.17[P17] was observed only in the zebrafish larvae model with a maximum log increase of 2.04 on 2 dpi. The HIE model did not support the replication of GII.17[P17].

Of the six GII.4 Sydney[P16] stool specimens tested, only four showed replication in both the HIE and zebrafish larvae models (Table 2. Interestingly, the two stool specimens that failed to replicate were not the same across both models. Each model failed to support replication of a different stool specimen. The two GII.4 Sydney[P16] stool specimens that replicated in both the models showed maximum log increases of 1.61 and 3.20 at 2 dpi in the zebrafish model. The corresponding log increases with the HIE model were 0.65 and 3.68 at 72 hpi, respectively. The two GII.4 Sydney[P16] strains that only replicated in the HIE model but not in the zebrafish model showed a log increase of 2.39 and 0.42 at 72 hpi. Whereas the two GII.4 Sydney[P16] strains that replicated in the zebrafish model exclusively had a maximum log increase of 2.56 at 3 dpi and 1.49 on 1 dpi.

## Discussion

In this study, the replication of seventeen different stool specimens comprised of three different GI HuNoV strains (GI.3[P3], GI.5[P5], and GI.7[P7]) and six GII strains (GII.3[P12], GII.4 Sydney[P16], GII.4 Sydney[P31], GII.4[P4 New Orleans], GII.6[P7], and GII.17[P17]) were compared using the zebrafish larvae and HIE models. Both models supported the replication of all five GI stool specimens tested. With GII stool specimens, 67% (8 out of 12) replicated in the zebrafish larvae model, and 58% (7 out of 12) replicated in the HIE model.

While the presence or absence of replication between different HuNoV strains can be compared between the HIE and zebrafish models, the level of replication cannot be directly compared due to fundamental differences in the experimental setup of each system. First, for the zebrafish larvae model, only a small inoculum volume (3 nL) of clarified stool specimen was microinjected into the yolk of each larva (Van Dycke et al., 2019), whereas in the HIE model, a much larger volume (100 μL) per well was used for infection (5). Secondly, different quantification methods were employed in this study for each of the models. For the zebrafish model, ddPCR was used to measure increase of viral RNA whereas for the HIE model RT-qPCR a standard curve of norovirus GI or GII transcript RNA to measure copy number increase was used.

Both the HIE and zebrafish larvae models have proven to be reliable platforms for studying HuNoV infectivity; however, they are limited by their inability to support replication of all HuNoV strains. For instance, in the present study, the HIE model did not support replication of the GII.17[P17] strain, whereas the zebrafish larvae model showed a significant 2 log increase in viral RNA copies post infection. Notably, the HIE model has previously been shown to support replication of GII.17 strains with P types P[38], P[13], and P[31] (6, 7). The P type is determined based on the RdRp region nucleotide diversity (17), and the RdRp region has been known to play a critical role in HuNoV replication (18). Previous studies have demonstrated that viral replication efficiency in the HIE model can vary depending on the intestinal segment of the donor and the strain specific compatibility of the HIE (8). Also, higher levels of replication of GII.3 strain has been observed in *STAT1* (master regulator of all three IFN (interferon) pathways and IFNAR1 (type I IFN receptor subunit) receptor knockout cell lines compared to wild type HIE cell lines showing that innate immunity affects replication of certain strains of HuNoV (10). It is possible that innate immune responses restricted its replication in this study. Six GII.4 Sydney[P16] specimens were evaluated across both models of which 67% (n=4) of the specimens replicated in both systems. This variability may be influenced by the presence of other substances in the stool specimens that affect replication efficiency. A previous study has shown stool composition to impact the stability and infectivity of astrovirus, another gastroenteritis virus (19). Stools have been shown to contain substances with potential antiviral activity such as innate immune factors like lysozyme and interferons, dietary compounds such as polyphenols, and microbial-derived compounds such as bacteriocins (19, 20). Interestingly, the two GII.4 Sydney[P16] specimens that replicated in the zebrafish model did not replicate in HIE, and vice versa. This suggests that, despite some similarities, differences in the ability to tolerate antiviral compounds present in the stool specimens by the models may contribute to the observed differences in replication.

Distinct differences in the expression of histo-blood group antigens exist between zebrafish and humans. Zebrafish possess five orthologues of human fucosyltransferase (*FUT*) genes (*fut*7 to *fut*11) which encode either α(1,3) or α(1,6)-fucosyltransferases. However, zebrafish do not have an orthologue of the human FUT2 gene and therefore lack α(1,2)-fucosyltransferases (21, 22, 23). In humans, *FUT*2 determines secretor status and individuals who lack functional *FUT*2 do not express ABH antigens in saliva and other bodily fluids (24). Notably, such non-secretor individuals are typically resistant to infection with many strains of HuNoV, as shown in human challenge studies (25, 26, 27, 28). Similarly, HIE cultures obtained from non-secretor individuals also do not support replication of GII.4, GII.17 and GI.1 (29). However, the absence of *FUT*2 gene in zebrafish did not limit their susceptibility to various HuNoV strains. This shows that zebrafish larvae express other receptors, which remain to be completely understood, that potentially aid in HuNoV entry and replication. Additionally, the HIE model required bile for GII.3 infection whereas the zebrafish larvae model did not require bile for replication.

In one study, different stool processing methods, such as serial filtration and serial centrifugation, significantly impacted the replication efficiency of HuNoV in various experimental culture models. The authors showed that serial filtration of stool specimens supported replication of HuNoV strains (n=3) in HIEs across different passage ages (passage 5 to 34), while serial centrifugation was associated with consistent replication only in newer HIE passage cultures (passage 5 to 19) (30). In zebrafish larvae models, centrifuged stool specimens are commonly used for microinjection (13, 14, 15). Although no direct comparisons have been made in the zebrafish model to assess how different stool processing methods affect replication, centrifugation retains vesicle associated HuNoV particles (31). In salivary gland cell lines, another model that has been reported to support replication of GII.4 Sydney strain, only vesicle containing HuNoV enriched inocula replicated efficiently (32). Similarly, for rotavirus, vesicle associated virions were more infectious than free viruses in mouse pups (31). In the present study, stool specimens were centrifuged for both the HIE model and the zebrafish model.

Most studies evaluating the replication of HuNoV strains using the HIE model have utilized enteroids derived from jejunal (J) cells (6, 5, 7, 8, 9, 10, 29). In 2024, HIE lines established from various segments of the small intestine (duodenum, jejunum, and ileum) obtained from four secretor positive individuals were compared for their ability to support replication of GII.3 and GII.4 strains. While HIEs from three donors supported replication of both strains, HIEs from one donor failed to support GII.3 replication but did support GII.4 (8). This highlights the influence of host genetic variability on HuNoV replication and suggests that the choice of HIE donor line can significantly affect experimental outcomes. In the current study, all HuNoV strains were tested using the J4FUT KI HIE line. Successful replication of specimens from older individuals as well as GI samples is likely because of the use of J4FUT2 KI line which was derived from a different (J4) donor. The J4FUT2 KI line has demonstrated increased susceptibility to HuNoV, supporting a broader range of GI and GII strains (8) and confirming that factors beyond just FUT2 expression may affect susceptibility. It is possible that certain strains which did not replicate in J4 might replicate in other HIE lines.

Initially, the zebrafish embryo model was considered, as a previous study demonstrated that embryos support higher replication efficiency of HuNoV and allow for easier microinjection (15) compared to the zebrafish larvae model which require anesthetizing the larvae before microinjection (13). However, in the present study, when embryos were microinjected with different stool specimens, they died within 1 to 4 h. This was likely due to the sensitivity of zebrafish embryos to toxic substances present in human stool specimens. As each stool specimens vary in composition, some may be less toxic to embryos, or if the viral titer is sufficiently high, dilution of the stool specimen may reduce toxicity without significantly lowering the virus load, thus allowing replication. This may explain why some HuNoV strains were able to replicate in zebrafish embryos without causing any mortality as reported by Tan and coauthors (2023)(15).

In conclusion, both the HIE and the zebrafish larvae model supported replication of most of the same stool samples containing GI and GII HuNoV strains. Notably, all GI positive samples tested replicated in both models. This comparative study suggests that zebrafish larvae express receptors similar to those in HIEs, to which various HuNoV strains can bind. Thus, these two models can be used in tandem to advance HuNoV research. However, each of the models offers distinct advantages and limitations. The HIE model closely mimics the human intestinal environment, including innate immune responses to pathogens, electron transport, and cell lifespan (33), but is costly and labor intensive. Additionally, the HIE model requires a larger volume of stool sample for inoculation. In contrast, the zebrafish larvae model is more affordable, scalable, suitable for high throughput screening, and requires low volume of stool sample for inoculation, though it may not fully replicate human immune responses (4). Also, zebrafish systems require appropriate housing setups which might not be available in all laboratories. Taken together, these models complement each other with regard to HuNoV pathogenesis and hence can be used together to accelerate the development of effective antiviral strategies to mitigate HuNoV transmission.

## Acknowledgements

This research was supported by an Arkansas Bioscience Institute grant awarded to K.E.G. from 2020 to 2023 and in part by the intramural food safety program and the Advanced Molecular Detection program at the Centers for Disease Control and Prevention (CDC). This research was also supported in part by the National Institute of Food and Agriculture (NIFA), U.S. Department of Agriculture (USDA) Hatch Act funding.

## Author Contributions

**Sahaana Chandran:** Conceptualization, Methodology, Investigation, Data curation, Writing – original draft, Writing – review & editing. **Jan Vinjé:** Conceptualization, Methodology, Writing – review & editing, Supervision, Project administration, Funding acquisition. **Kimberly Huynh:** Investigation, Data curation, Writing – review & editing. **Kristen E. Gibson:** Conceptualization, Methodology, Writing – review & editing, Supervision, Project administration, Funding acquisition.

